# Decoding music-evoked emotions in the auditory and motor cortex

**DOI:** 10.1101/2020.05.24.101667

**Authors:** Vesa Putkinen, Sanaz Nazari-Farsani, Kerttu Seppälä, Tomi Karjalainen, Lihua Sun, Henry K. Karlsson, Matthew Hudson, Timo T. Heikkilä, Jussi Hirvonen, Lauri Nummenmaa

**Affiliations:** Turku PET Centre, and Turku University Hospital, University of Turku, Finland; Department of Radiology, Turku University Hospital, Finland; Department of Psychology, University of Turku, Finland

**Keywords:** Music, Emotion, fMRI, MVPA, pattern-classification

## Abstract

Music can induce strong subjective experience of emotions, but it is debated whether these responses engage the same neural circuits as emotions elicited by biologically significant events. We examined the functional neural basis of music-induced emotions in a large sample (n=102) of subjects who listened to emotionally engaging (happy, sad, fearful, and tender) pieces of instrumental music while their haemodynamic brain activity was measured with functional magnetic resonance imaging (fMRI). Ratings of the four categorical emotions and liking were used to predict haemodynamic responses in general linear model (GLM) analysis of the fMRI data. Multivariate pattern analysis (MVPA) was used to reveal discrete neural signatures of the four categories of music-induced emotions. To map neural circuits governing non-musical emotions, the subjects were scanned while viewing short emotionally evocative film clips. The GLM revealed that most emotions were associated with activity in the auditory, somatosensory and motor cortices, cingulate gyrus, insula, and precuneus. Fear and liking also engaged the amygdala. In contrast, the film clips strongly activated limbic and cortical regions implicated in emotional processing. MVPA revealed that activity in the auditory cortex in particular as well as in the primary motor cortices reliably discriminated the emotion categories. Our results indicate that different music-induced basic emotions have distinct representations in regions supporting auditory processing, motor control, somatosensation and interoception but do not strongly rely on limbic and medial prefrontal regions critical for emotions with survival value.

## 1. Introduction

Human attraction to music stems from the aesthetic experience and strong emotional reactions music evokes. Basic emotion theories posit a finite number of innate and universal emotions that have evolved because they promote survival by motivating adaptive behaviors and help maintaining physiological homeostasis (Ekman, 1992), and numerous neuroimaging studies have tested whether music is capable of activating the survival circuits underlying these emotions (Brattico et al., 2011; Koelsch et al., 2013; Mitterschiffthaler et al., 2007). Indeed, emotional responses to music show many hallmarks of basic emotions including distinct subjective experience, emotional expression, action tendencies, changes in the autonomic nervous system activation (Juslin & Vastfjall, 2008) and distinct effects of thought and judgement (Vuoskoski & Eerola, 2012). Yet, it seems counterintuitive that that instrumental music could elicit “genuine” basic emotions because emotional reactions to music do not have unambiguous causes (real-life threat, success, or loss etc.) and they do not involve events that have obvious goal or survival-relevant consequences (Konečni, 2008; Scherer & Zentner, 2008). Thus, even though listeners often interpret their subjective reactions to music as basic emotions, these responses may differ from genuine basic emotions in the underlying neural processes.

Some acoustic features contributing to different music-induced emotions may be universally recognizable (Fritz et al., 2009), perhaps because similar cues communicate emotion in human vocalizations and music (Frühholz et al., 2016; Juslin & Laukka, 2003). These features may recruit partly the same neural networks across the domains (Escoffier et al., 2013; Peretz, 2010). Music-induced emotions are nevertheless dependent on cultural learning (Higgins, 2012; McDermott et al., 2016) and therefore may engage partly distinct neural systems than the putatively innate basic emotions.

Emotions elicited by biologically salient stimuli consistently engage subcortical structures such as the amygdala, thalamus, striatum, and brain stem and cortical regions such as the anterior cingulate, insula, and medial and orbital frontal cortex (Kober et al., 2008; Vega et al., 2016). A recent meta-analysis indicates that different music-induced emotions also activate these structures suggesting that these emotions may rely on some of the same evolutionarily adaptive mechanisms that underlie emotions with survival value (Koelsch, 2014). However, it is unclear whether the activation of these circuits discriminates between specific categories of music-induced emotions. Thus far, fMRI studies have largely failed to find reliable category-specific correlates of music-induced emotions and often contradict each other in terms of the regions they implicate in specific music-induced emotions such as happiness (Brattico et al., 2011; Mitterschiffthaler et al., 2007), sadness (Mitterschiffthaler et al., 2007; Trost et al., 2012) or fear (Bogert et al., 2016; Koelsch et al., 2013).

Mass univariate general linear model (GLM) has been the dominating analysis approach in fMRI studies on emotions. More recently, multivariate pattern analysis (MVPA) has been adopted in order to decode of high-dimensional regularities in fMRI data that predict different basic and social emotions (Nummenmaa & Saarimäki, 2017). These studies indicate that discrete emotions perceived in facial expressions, bodily movements and vocalizations (Ethofer et al., 2009; Peelen et al., 2010) can be decoded from brain activity in sensory cortices while emotional states induced by stories, movies and imagery are represented in discrete and widely distributed activity patterns across various cortical and subcortical regions (Kassam et al., 2013; Saarimäki et al., 2015; Sitaram et al., 2011). Thus, different music-evoked emotions might also be based on distinct but spatially overlapping neural activation patterns not captured by univariate analysis. Two prior MVPA-fMRI studies indicate that activity patterns in the auditory cortices predict whether subjects heard short bursts of violin and clarinet sounds conveying either sadness, happiness or fear (Sébastien Paquette et al., 2018; Sachs et al., 2018). These activity patterns probably reflect sensory processing of acoustic properties with different emotional connotations, but it is unclear whether such auditory cortical patterns predict different emotions actually felt by the listeners during naturalistic music-listening. A few other emotion-related MVPA-fMRI studies have used music as stimuli but either collapsed data obtained with music and other means of emotion-induction precluding inferences specifically about music-induced emotions (Kragel & LaBar, 2015) or focused only on emotional valence instead of discrete music-induced emotions (Kim et al., 2017). Thus, it remains unclear whether distinct music-induced emotions are represented in anatomically distributed activity patterns similarly to emotions elicited by non-musical stimuli.

### The current study

Here we combined GLM and multivariate pattern analyses of fMRI data from a large sample of 102 subjects to test whether i) different music-induced emotions reliably activate brain’s emotion circuit similarly as emotions elicited by biologically salient episodes shown in videos, and ii) whether music-evoked emotional responses are organized in a discrete fashion in the brain. We focused on emotions happiness, sadness, fear, and tenderness because they cover the four quadrants of the valence-arousal-circumplex, are included in categorical models frequently used to describe music-induced emotions in the literature (Eerola & Vuoskoski, 2010; Zentner et al., 2008), and are rated consistently across listeners (Juslin, 2013). Subjects underwent fMRI scanning while they listened to excerpts of instrumental movie soundtracks validated previously to reliably elicit these emotions. To map neural circuits governing non-musical emotions, we also scanned the same subjects in a previously validated emotion “localizer” paradigm (Karjalainen et al., 2017; Lahnakoski et al., 2012) in which subject view film clips with positive and negative emotional content. Like music, fictious films are devoid of real-life consequences and yet elicit strong emotions. However, unlike instrumental music, movies offer a rich, multimodal simulation of the real-life emotional events. Thereby, movies provide a naturalistic paradigm for mapping neural circuits supporting emotions in real-world situations (Sonkusare et al., 2019). We found that music-induced basic emotions engaged regions supporting auditory processing, motor control, somatosensation and interoception but did not strongly activate limbic and medial prefrontal regions that govern non-musical emotions. Furthermore, MVPA revealed that activity in the auditory and primary motor cortices, but not in the core emotion circuits, reliably discriminated the emotion categories.

## 2. Materials and methods

### Subjects

Altogether 102 volunteers participated in the study (51 females, mean age 31 years, range 20–57 years). Fifty percent of subjects had received no formal training on a musical instrument (The median for the months of training for the was 6.3). The exclusion criteria included a history of neurological or psychiatric disorders, alcohol or substance abuse, current use of medication affecting the central nervous system and the standard MRI exclusion criteria. Two additional subjects were scanned but excluded from further analyses because unusable MRI data due to gradient coil malfunction. All subjects gave an informed, written consent and were compensated for their participation. The ethics board of the Hospital District of Southwest Finland had approved the protocol and the study was conducted in accordance with the Declaration of Helsinki.

### Study design

The stimuli were eighteen 45-sec (including a 10-ms fadeout) excerpts of instrumental music. Sixteen excerpts were chosen from a set of movie soundtracks based on high ratings for happiness, sadness, fear or tenderness in a previous study (Eerola & Vuoskoski, 2010). For each of the four emotions, four stimuli were selected that had received high rating for the target emotion and low ratings for the other three (see the supplementary material for details). Two additional excerpts of instrumental rock (Far Beyond the Sun by Yngwie J. Malmsteen) were included to add variation in the musical material in terms of genre and instrumentation. Instrumental music was used to minimize the contribution of semantic knowledge on the emotional responses.

A 45-sec random tone sequence followed by 45 seconds of silence was presented at the beginning of the run. After this, the musical excerpts were presented in a fixed pseudo-random order without silent breaks in between. The last excerpt was followed by 45 seconds of silence, another 45-sec random tone sequence and a third 45-sec silent period (see the Supplementary Material for the contrasts between music vs. control stimuli and music vs. silence). Subjects were asked to remain still during the fMRI and focus on the feelings evoked by the music. Stimuli were presented binaurally via MRI-compatible headphones (Sensimetrics S14) at a comfortable level adjusted individually for each participant. After the scanning session, the subjects listened each music excerpt again and rated for experience of fear, happiness, sadness, tenderness, and liking on a scale ranging from 1-10 (1 = extremely weak, 10 = extremely strong). Ratings were done using an online rating tool developed in-house (https://gitlab.utu.fi/tithei/pet-rating). Complete ratings were obtained from 91 participants. Ratings averaged across these subjects were used as regressors in the analysis of the BOLD data (cf. Trost, et al., 2012).

To map brain regions governing non-musical emotions, we used a previously validated emotion “localizer” paradigm in which the subjects viewed a compilation of 96 movie clips extracted from mainstream English language feature films (mean duration 12.5 s; total duration 20 minutes) containing variable emotional and non-emotional content (for details, see Karjalainen et al., 2017, 2019; Lahnakoski et al., 2012). The movie clips were presented in fixed order without breaks in between. The film clips contained scenes with displays of positive emotions (e.g. laughter, friendly discussion, expression of affection between parents and children or between romantic partners), negative emotions (e.g. crying, arguing, violence) as well as scenes without emotional content (e.g. humans in a neutral emotional state, objects, landscapes). Dynamic ratings with a 4 sec temporal resolution were obtained for the intensity of positive and negative emotions observed in the film clips from a separate sample of subjects (n=6) who did not participate in fMRI study. The average ratings were subsequently used as regressors in GLM analysis. The film clips were presented via NordicNeuroLab VisualSystem binocular display (audio was delivered as described above).

### MRI data acquisition

The MRI data were acquired using a Phillips Ingenuity TF PET/MR 3T whole-body scanner. High-resolution (1 mm^3^) structural images were obtained with a T1-weighted sequence (TR 9.8 ms, TE 4.6 ms, flip angle 7°, 250 mm FOV, 256 × 256 reconstruction matrix). 407 functional volumes were acquired with a T2*-weighted echo-planar imaging sequence (TR 2600 ms, TE 30 ms, 75° flip angle, 240 mm FOV, 80 × 80 reconstruction matrix, 62.5 kHz bandwidth, 3.0 mm slice thickness, 45 interleaved slices acquired in ascending order without gaps).

### Structural and functional MRI data preprocessing

MRI data were preprocessed using fMRIPprep 1.3.0.2 (Esteban et al., 2019). The following preprocessing was performed on the anatomical T1-weighted (T1w) reference image: correction for intensity non-uniformity, skull-stripping, brain surface reconstruction, spatial normalization to the ICBM 152 Nonlinear Asymmetrical template version 2009c (Fonov et al., 2009) using nonlinear registration with antsRegistration (ANTs 2.2.0) and brain tissue segmentation. The following preprocessing was performed on the functional data: co-registration to the T1w reference, slice-time correction, spatial smoothing with an 6mm Gaussian kernel, automatic removal of motion artifacts using ICA-AROMA (Pruim et al. 2015) and resampling the MNI152NLin2009cAsym standard space. Low-frequency drifts were removed with a 240-s- Savitzky–Golay filter (Çukur et al., 2013).

### Full-volume GLM data analysis

The fMRI data analyzed in SPM12 (Wellcome Trust Center for Imaging, London, UK, (http://www.fil.ion.ucl.ac.uk/spm). To reveal regions activated by music vs. silence, a general linear model (GLM) was fitted to the data where the music sequence and the silent periods were modelled with separate boxcar regressors. The regional effects of the five emotion dimensions were assessed with a GLM where the design matrix included the music boxcar regressor and each of the emotion rating regressors as parametric modulators. Contrast images for the main effects of music, silence and each emotion regressor were generated for each participant and subjected to a second-level analysis. Clusters surviving family wise error (FWE) correction (P < 0.05) are reported.

### Region-of-Interest Analyses

Beta weights for each of the five emotion regressors were extracted from 15 anatomical regions-of-interest (ROI) implicated in emotional processing (Sébastien Paquette et al., 2018; Saarimäki et al., 2015, 2018). These included the amygdala, caudate, putamen, pallidum, thalamus, hippocampus, insula, anterior and posterior cingulate, SMA, precentral and postcentral gyri, precuneus, frontal pole, and auditory cortex. The mean beta weights for each ROI were calculated from the first-level contrast images of each subject using ROI masks derived from the AAL atlas. The beta weights for each emotion in each ROI were compared against zero with separate one-sample t-tests.

### Multivariate pattern analysis

A between-subject classification of the four emotion categories (fear, happiness, sadness, and tenderness) was performed in Python using the PyMVPA toolbox (Hanke et al., 2009). A support vector machine with radial basis function (RBF) kernel was trained to recognize the emotion categories using leave-one-subject-out cross-validation, where the classifier is trained on the data from all except one subject and tested on the hold-out subject data; this procedure is repeated N times so that each subject is used once as the hold-out subject; this kind of leave-one-subject-out cross-validation tests the generalizability of the results across the sample of the subjects. A subject-wise GLM with regressors for each stimulus was fit to the data resulting in 16 beta weights (4 emotion categories × 4 song per category) per voxel for each subject and the MVPA was applied to the beta images The beta weights were normalized to have zero mean and unit variance before application of MVPA. This analysis was performed using whole-brain data (with non-brain voxel masked out) and within a subset of the ROIs (amygdala, hippocampus, thalamus, anterior and posterior cingulate, SMA, precentral and postcentral gyri, precuneus, frontal pole, auditory cortex) where emotion classification has been successful in previous studies (Sébastien Paquette et al., 2018; Saarimäki et al., 2015). In the whole-brain MVPA, an ANOVA feature selection was applied to the training set within each cross-validation where 5000 voxels with the highest F-score were selected. Classification accuracy was quantified by computing the proportion of correctly classified songs relative to the number of songs in each of the four categories (i.e. recall). To estimate the null distribution for the classification accuracies (naïve chance level 25%) the following procedure was repeated 500 times: 1) we randomly shuffled emotion category labels, 2) ran the whole-brain MVPA with 102 leave-one-subject-out cross-validations, where the classifier was trained on the data with shuffled labels from N-1 subject and tested on data with correct labels from the remaining subject, and 3) calculated the classification accuracies on each of the 500 iterations. If the true accuracy was larger than 95 percent of the accuracies obtained with the randomly shuffled labels, the true accuracy was considered significant with an alpha of 0.05

## 3. Results

### Behavioral Ratings

The emotion ratings indicated that the musical excerpts reliably induced the target emotions (**Figure 1 a**), and most pieces elicited moderate feelings of liking (mean: 5.7, **supplementary figure 1**). The pieces were rated consistently across the subjects for fear, happiness, sadness, tenderness and liking (mean inter-subject correlation: .68). Repeated measures ANOVA’s revealed significant main effects of song category on the mean ratings for each emotion (*p* < .001, for all emotions, see supplementary materials for the full ANOVA results). Pair-wise comparisons of fear, sadness, happiness and tenderness ratings across the scary, sad, happy, and tender songs showed that each song category was rated highest for the target emotion (i.e. the scary songs were rated higher for fear than the other songs etc., *p* < .001 for all contrasts). Liking ratings were lower for the scary songs than for any other song category (*p* < .05 for all comparisons) and lower for the rock songs than for the happy songs (*p* < .01). We computed dissimilarity matrices (Euclidean distances) for each song pair to illustrate the similarity/dissimilarity of rating profiles within and across the emotion categories (**Figure 1 b**). The dissimilarity matrix indicates that the rating profiles were more similar within than across categories, confirming that the emotional experiences were similar for songs within a given category and distinct from the experience induced by the songs in the other categories.

**Figure 1.**
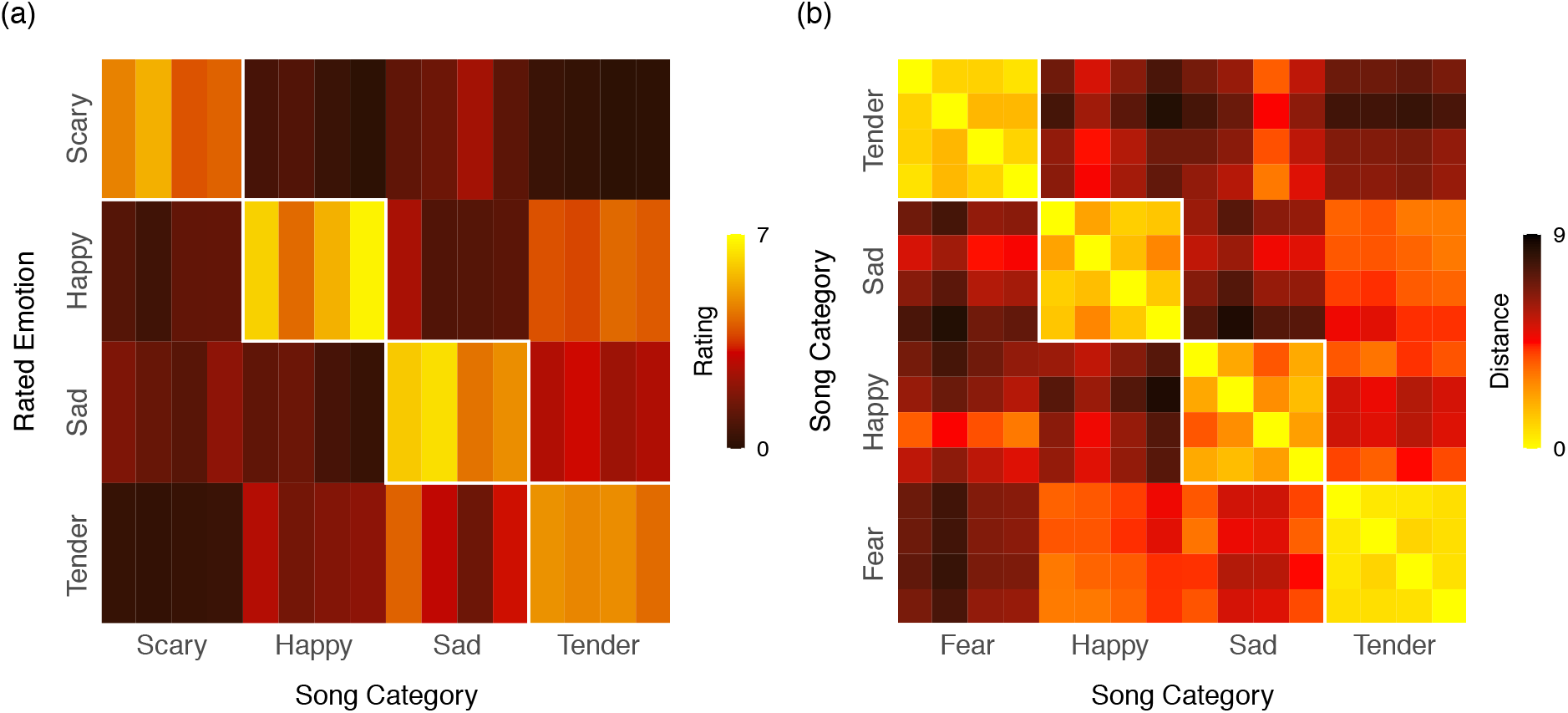
(a) Mean ratings for the intensity of each emotion for each musical excerpt. (b) Rating dissimilarity matrix (Euclidean distance) for each song pair.

### Brain responses evoked by emotional music

We first modelled brain responses to each musical emotion dimension. **Fear** elicited subcortical activity bilaterally in the brainstem, thalamus, putamen and pallidum. Activity was also observed in the amygdala and the insula (**Figure 2**). There was extensive bilateral activity across the precentral gyrus and SMA extending into the postcentral gyrus. A cluster of activity was also found in the cerebellum. Frontal activity was found in the right inferior frontal gyrus and bilaterally in frontal operculum and frontal pole. The cingulate gyrus was activated across both anterior and posterior portions. Occipital activity was observed in the precuneus. Finally, there was activity across the auditory cortical regions including Heschl’s gyrus, planum temporale, planum polare and suprerior temporal gurys. **Happiness** elicited right-lateralized activity in the pallidum and putamen. The insula was activated bilaterally. Parietal activity was observed in the SMA. Frontal activity was found in the right frontal pole. There were significant clusters of activity in the anterior and posterior divisions of the cingulate gyrus. Occipital activity was observed in the cuneus and precuneus. Finally, there was activity in Heschl’s gyrus extending into planum temporale**. Sadness** activated the insula and frontal operculum, anterior cingulate gyrus, cuneus/precuneus, and Heschl’s gyrus. **Tenderness** elicited subcortical activity in the thalamus, pallidum and the caudate. Occipital activity was found in the right precentral gyrus. Frontal activity was observed in inferior, middle and superior frontal gyri and the frontal pole. Significant clusters were also found bilaterally in the supramarginal gyrus. Finally, there was extensive activity in the auditory cortical regions encompassing Heschl’s gyrus, planum temporale, planum polare and superior temporal gyrus. **Liking** activated the brainstem, hippocampus and parahippocampal gyrus, amygdala, putamen, and thalamus. Extensive activity was observed across the somatosensory and motor regions in the precentral and postcentral gyri, SMA and cerebellum. Frontal activity was observed bilaterally in the inferior, middle and superior frontal gyri and frontal pole. Occipital activity was seen in the lingual gyrus. Significant clusters were also found bilaterally in the supramarginal gyrus and the superior temporal gyrus. To assess which regions were most consistently activated by different emotions, we first binarized the aforementioned statistically thresholded activation maps for each emotion (fear, happiness, sadness, tenderness) and generated a summed image of the maps. The resulting cumulative activation map (**Figure 3A**) thus shows how many emotion categories activated each voxel at the a priori (FWE 0.05) statistical threshold.

**Figure 2.**
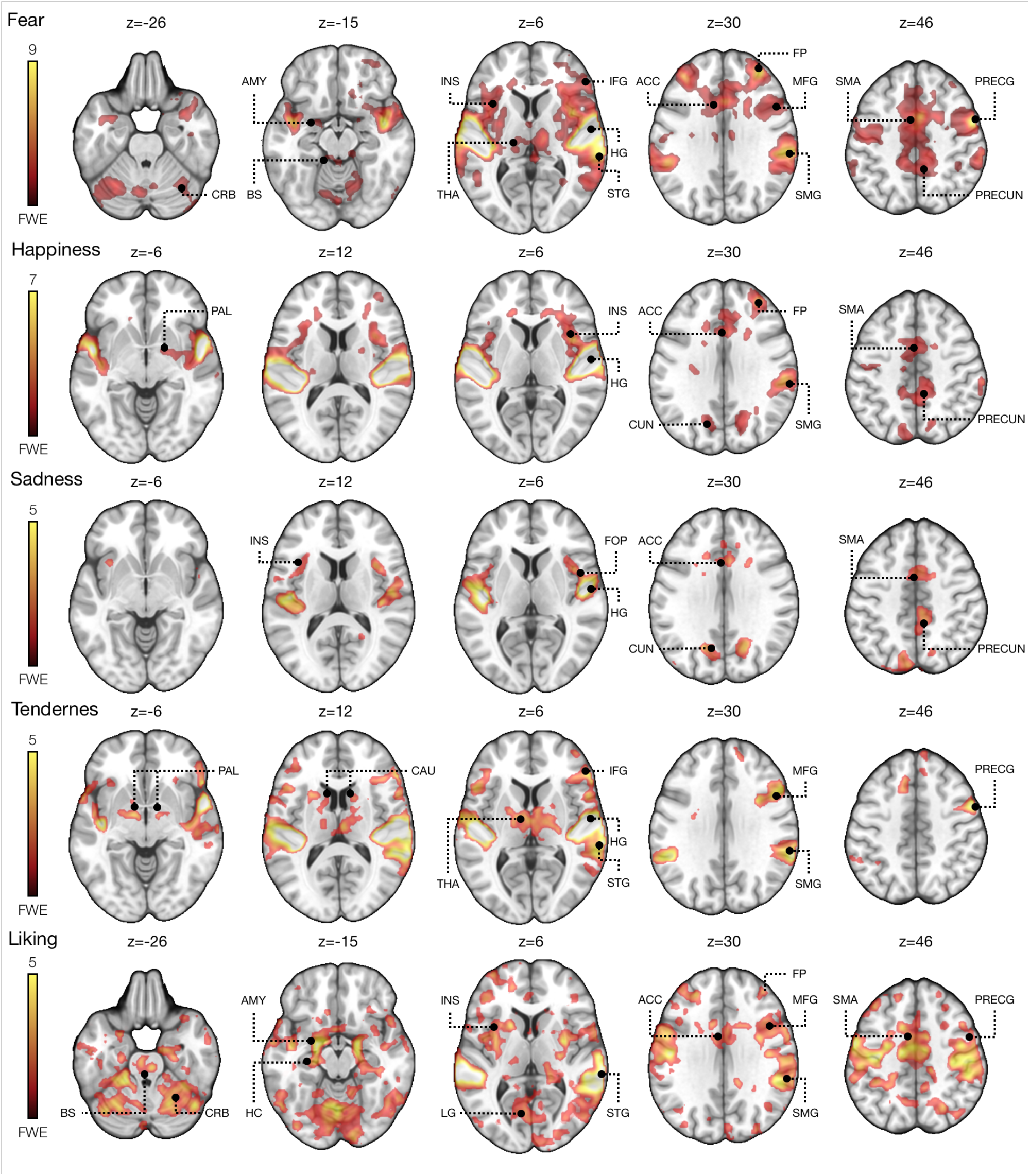
Brain regions responding to fear, happiness, sadness, tenderness and liking. The data are thresholded at *p* < .05 and FWE corrected at cluster level. ACC = Anterior Cingulate, AMY = Amygdala, CAU = Caudate, CBR = Cerebellum, CUN = Cuneus, FOP = Frontal Operculum, FP = Frontal Pole, HC = Hippocampus, HG = Heschl’s Gyrus, IFG = Inferior Frontal Gyrus, INS = Insula, LG = Lingual Gyrus, MFG =Middle Frontal Gyrus, PAL = Pallidum, PRECG = Precentral gyrus, PRECUN = Precuneus, SMA = Supplementary Motor Area, SMG = Supramarginal gyrus, STG = Superior Temporal Gyrus, THA = Thalamus. The color bar indicates T-value.

**Figure 3.**
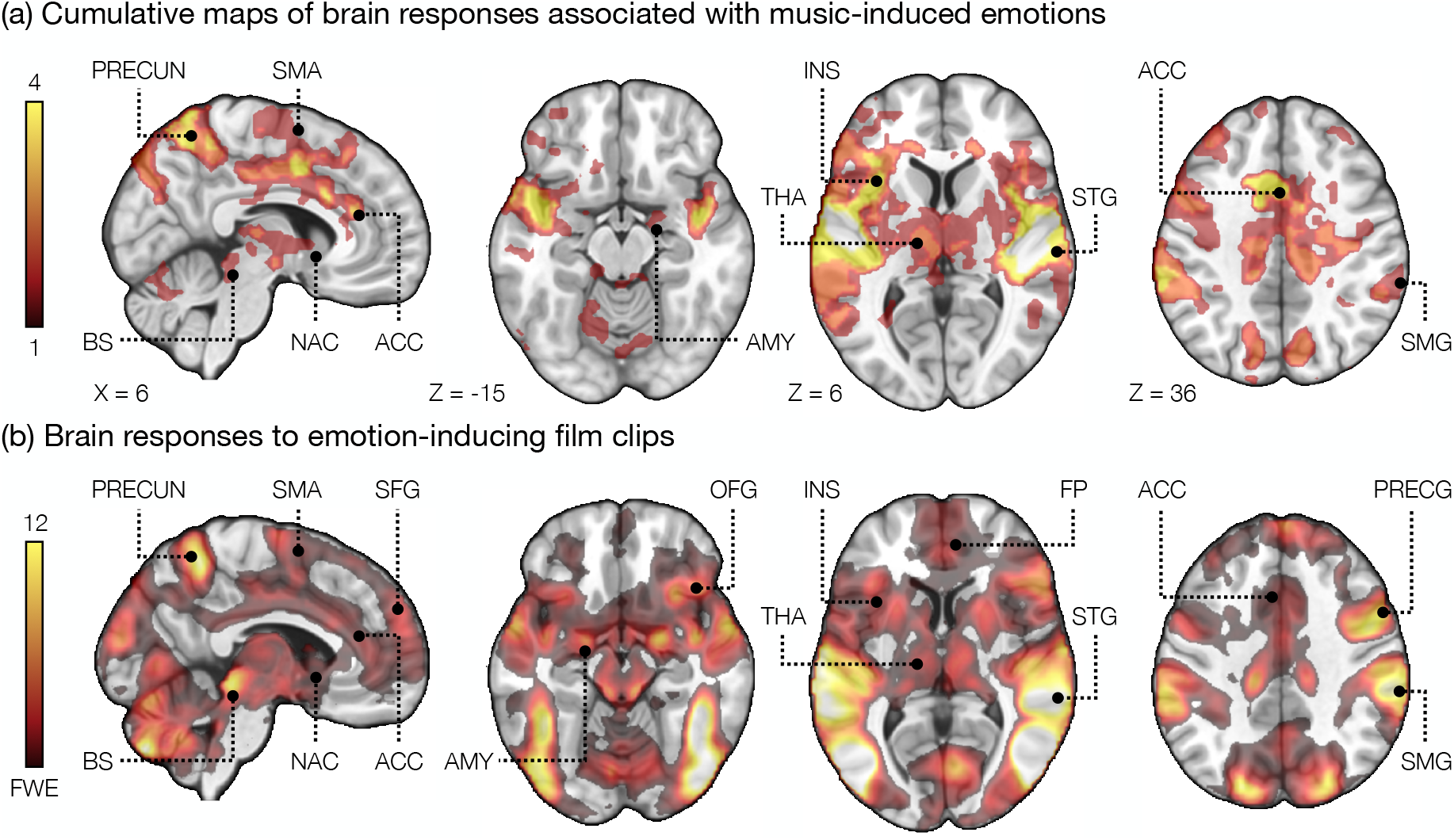
(a) Cumulative regional map of music-evoked activations. Voxel intensities show the number of musical categories (across fear, happiness, sadness and tenderness) showing statistically significant (p < 0.05 FWE) activations at each voxel (b) Brain regions responding to emotional stimulation in the film experiment. The data are thresholded at *p* < .05 and FWE corrected at cluster level. ACC = Anterior Cingulate, AMY = Amygdala, FP = Frontal Pole, INS = Insula, PRECG = Precentral gyrus, PRECUN = Precuneus, SMA = Supplementary Motor Area, STG = Superior Temporal Gyrus, THA = Thalamus. The color bar indicates T-value.

### Brain responses to emotional videos

In the GLM analysis of the video experiment data, the emotion intensity regressor revealed activation in various regions associated with emotion such as the brainstem, thalamus, ventral striatum (nucleus accumbens, NAcc), amygdala, insula and the cingulate gyrus and the orbitofrontal cortex (Figure 3b). There was also extensive activation in the midline frontal regions and across somatomotor regions in the precentral gyrus, SMA and the cerebellum. Occipital activity was observed in the precuneus and in regions involved in visual processing. Temporal activity was observed in the auditory cortical regions

### Regional effects to emotional music

To parse out the emotion-specificity of the regional responses, we next performed ROI analyses for the music-evoked responses (Figure 4). Only fear and liking significantly activated the amygdala, putamen and the ACC at the ROI level. Furthermore, significant hippocampal activation observed only for liking. Happiness activated the insula which was also activated by fear. Tenderness activated the thalamus and pallidum. Sadness did not reveal any significant effects in the analyzed ROIs, and no effects were observed in the caudate.

**Figure 4.**
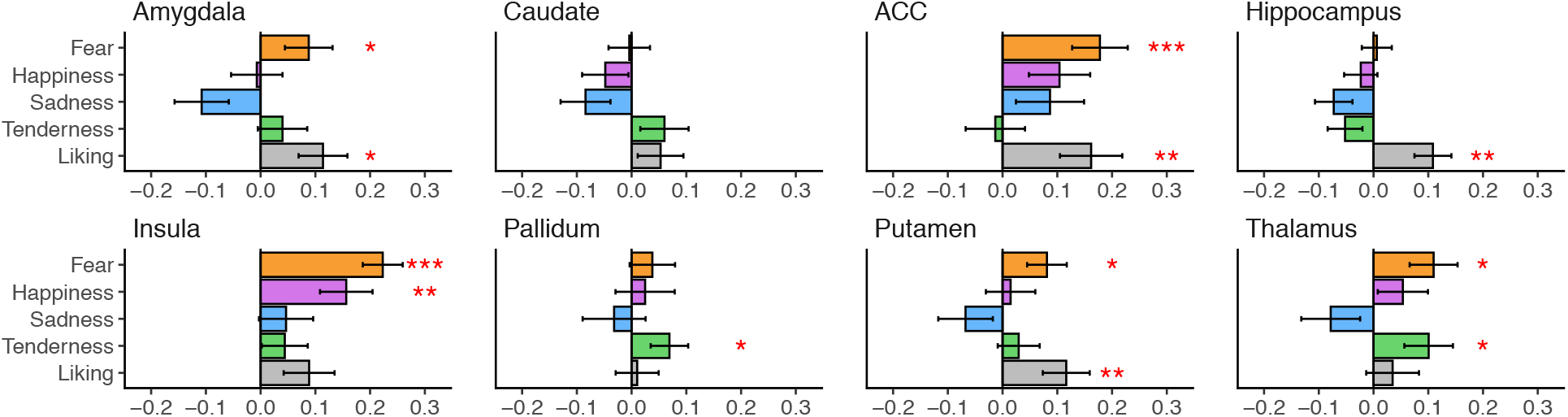
Mean beta weights (arbitrary scale) for each emotion regressor in each ROI. The error bars indicate standard error of mean and asterisks denote significant one sample t-tests against zero. The regional results are plotted for visualization purposes only, statistical inference is based on the full-volume analysis. * p < .05, ** p <.01, *** p<.001

### Emotion classification with MVPA

For the whole-brain four-class MVPA, the mean accuracy was 49% and each of the four emotions were classified significantly above chance level (fear: 51%, happiness: 45%, sadness: 53%, tenderness: 48%, all *p:*s < .001). The 5000 most important voxels for the multi-class classification were predominantly located in the bilateral auditory cortices (**Figure 5a**). Accordingly, the ROI-level MVPA revealed that classification accuracy for all four emotions was clearly above chance level for the auditory cortex (mean accuracy 48%) and reached significance also in the precentral gyrus (mean accuracy 29 %) (**Figure 5 b, figure 6**). Although the overall classification accuracy significantly exceeded the chance level for the amygdala, thalamus, precentral and postcentral gyri, SMA and precuneus ROIS, the classification accuracies were dramatically lower than that of the auditory cortex. Furthermore, none of these ROIs significantly discriminated between all four emotions (**Figure 5 b, supplementary Figure 5**). The ROI analyses were also performed separately for the left and right hemispheres but these analyses revealed no substantial hemispheric differences and the classification accuracies tended to be lower than for the bilateral ROI analyses (see supplementary Figure 5).

**Figure 5.**
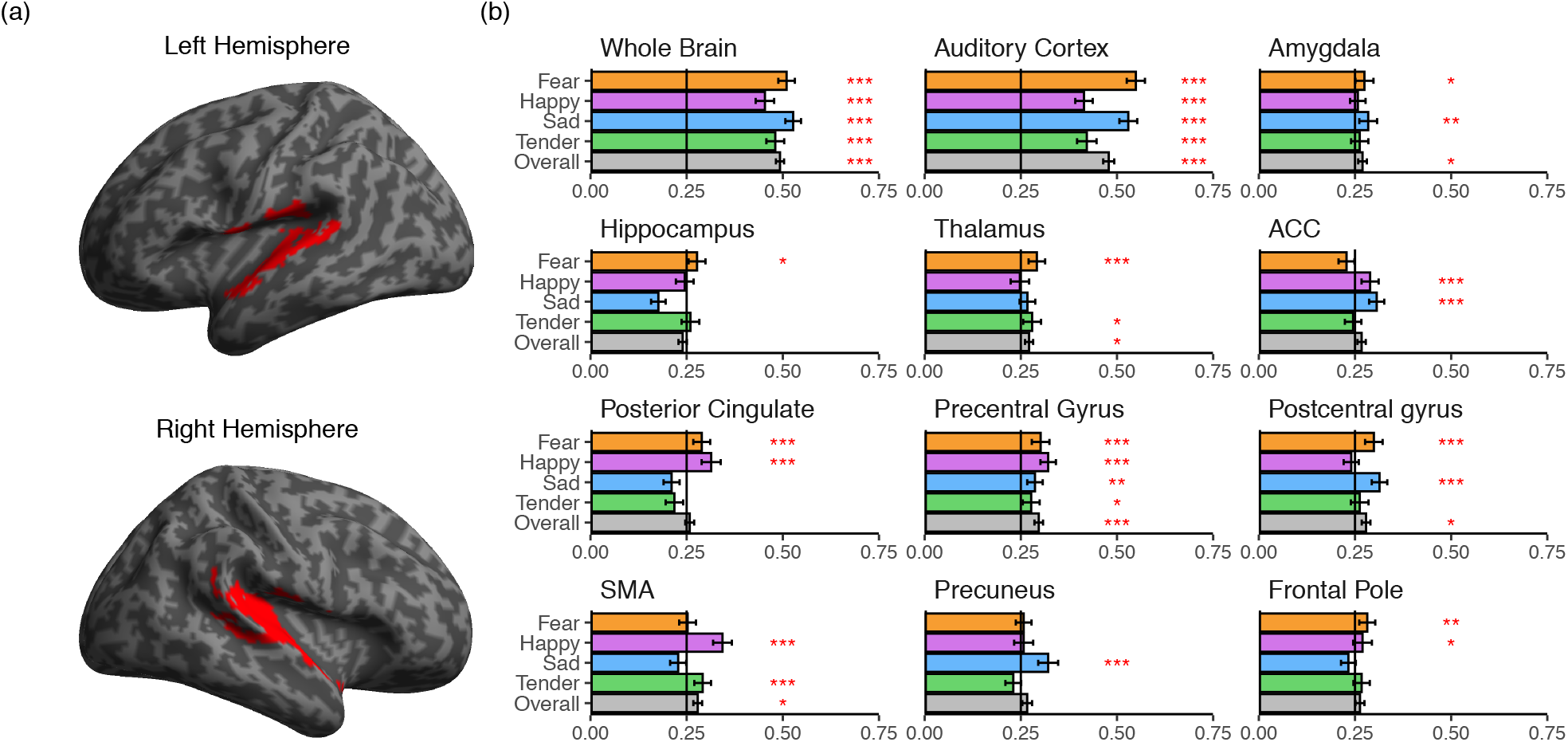
(a) The most important voxels for the between-subject classification of emotion categories in the whole-brain analysis. (b) Emotion-wise classification accuracies for the whole-brain and regional MVPA. * p < .05, ** p <.01, *** p<.001.

**Figure 6.**
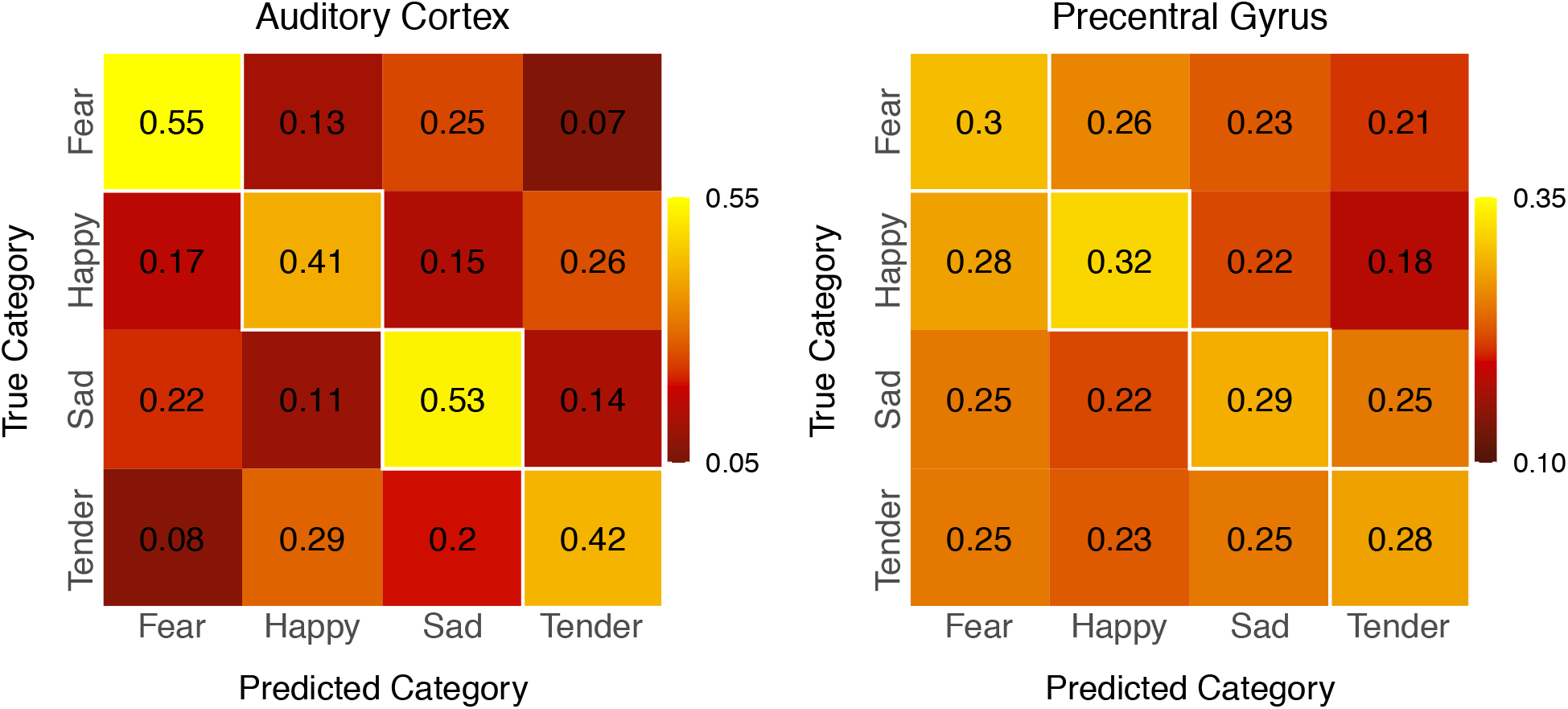
MVPA confusion matrices for the auditory cortex and precentral gyrus ROIs. The numbers indicate the proportion of true positives (diagonal) and false negatives within each emotion category.

## 4. Discussion

Our main finding was that music-induced fear, happiness, sadness and tenderness have discrete neural representations in the auditory and motor cortices. The GLM results showed that the music-induced emotions activated a common network comprising the auditory cortices, somatosensory/motor regions (SMA, precentral and postcentral gyri, cerebellum), cingulate cortex, insula, and precuneus, yet classification accuracies did not consistently exceed chance level in any region outside auditory and motor cortices. Moreover, music-induced emotions resulted in weak activation in limbic regions such as the ventral striatum, thalamus and amygdala and cortical areas such as medial prefrontal and orbitofrontal cortices which are implicated in non-musical emotions (Lindquist et al., 2012) and were strongly activated by the emotional film clips. These findings cast doubt on the argument that different music-induced emotions engage the affective circuits in a discrete fashion (c.f. Saarimäki et al 2015; 2018) and instead suggest that these emotions involve a relatively unspecific activation outside the auditory and motor cortices.

### Auditory and motor cortical representations of music-induced basic emotions

Despite high statistical power and subjective reports of strong music-evoked emotion, we found only limited evidence that music-induced emotions consistently engage the same core limbic circuits as emotions with survival value. The GLM analysis revealed extensive auditory cortical activation for all four emotions as well as liking. This accords with previous studies showing that the auditory cortices are more strongly activated by happy and sad than neutral music (Mitterschiffthaler et al., 2007) and that hemodynamic response in the auditory cortex tracks dynamic arousal ratings during music listening (Trost et al., 2012) and when listening to speech describing emotional episodes with neutral prosody (Nummenmaa et al., 2014). This effect likely reflects enhanced, general sensory gain for the emotionally salient stimuli. In the whole-brain MVPA, activity patterns in the auditory cortices also predicted with high accuracy whether the subjects were listening to a scary, happy, sad or tender piece indicating that the BOLD signal in bilateral auditory cortices carries sufficient information for distinguishing categories of musical pieces conveying different discrete emotions. The classification accuracies we observed in the auditory cortex ROIs are comparable with those reported in prior pattern classification studies on musical (Paquette et al., 2018; Sachs, et al., 2018) and non-musical emotions (Saarimäki et al., 2016). Tonal, rhythmic and timbral differences between sad, happy, scary and tender music (Laurier et al., 2009) presumably contribute to these emotion-specific auditory cortical activity patterns. These patterns were consistent across individuals, as evidenced by the successful leave-one-subject-out cross-validation. In the regional level MVPA, the auditory cortex showed high classification accuracy for all emotions, whereas the accuracies in all other ROIS constituting central nodes of the emotion circuit were substantially lower or at chance-level.

For the whole-brain and auditory cortex ROI, the classification accuracies for happiness and tenderness were slightly lower than for fear and happiness. The auditory cortex confusion matrix shows that these categories were somewhat more difficult to distinguish from one another than the other categories. Furthermore, the rating dissimilarity matrix suggests that the emotional experience was slightly more similar between happy and tender pieces (mean Euclidean distance: 3.82) relative to the other categories (mean Euclidean distance across other category pairs: 6.01). Thus, the similarity in emotional experience was reflected in similarity of the corresponding brain activity patterns in the auditory cortex. This probably reflects the fact that both categories included pleasant sounding pieces in major key that received high liking ratings.

Primary motor cortex (precentral gyrus) was the only ROI outside the auditory cortex where classification accuracy exceeded the chance level for all the tested emotions. The SMA and postcentral gyrus ROIs also showed above chance-level overall classification accuracy, although the accuracies for all individual emotions did not consistently reach significance in these ROIs. In the GLM most emotions also activated regions involved in motor control and somatosensation such as the precentral and postcentral gyri, SMA, supramarginal gyrus and cerebellum. This accords with data implicating somatomotor regions in emotion perception and the subjective experience of emotions (Adolphs et al., 2000; Nummenmaa et al., 2012; Pourtois et al., 2004) and studies indicating that pleasurable music engages motor regions such as the SMA and cerebellum more strongly than emotionally less evocative music (Blood & Zatorre, 2001; Pereira et al., 2011). Music often induces spontaneous rhythmic movements in the listener which is evident already in young children (Zentner & Eerola, 2010) and even in some non-human animals (Hattori & Tomonaga, 2020). Thus, the sensorimotor activity observed in the current study might reflect this perception-movement coupling even in the absence of overt movement (cf. Zatorre, Chen, & Penhune, 2007). Furthermore, different music-induced emotions prime distinct types of movements (Burger et al., 2013) which could explain the covariation of activity in the motor regions and emotion ratings in the GLM and the emotion specific multivoxel activity patterns in primary motor cortex.

Two prior MVPA-fMRI studies (Sébastien Paquette et al., 2018; Sachs et al., 2018) have found evidence for differential activity patterns for fear, happiness and sadness in the auditory cortex and importantly, neither of these could reveal emotion-specific activity pattern within the limbic regions. Arguably, these results might stem from use of short (<2 sec) sound bursts, from the ‘Musical Emotional Bursts’ stimulus set (Paquette et al., 2013)as stimuli which probably did not evoke strong subjective emotions in the subjects (although neither of these studies report the intensity of the felt emotions in response to the stimuli but only accuracy of behavioral classification to the a priori emotional categories). However, despite i) using naturalistic music as stimuli rated relatively high on each target emotion and ii) having high statistical power with 102 subjects, the present categorical approach provided no evidence of emotion-specific activation outside the auditory and motor cortices. This is in clear contrast with pattern recognition studies in other domains which have found that emotion categories are represented in a distributed fashion across cortical and subcortical regions (Kragel & LaBar, 2016; Nummenmaa & Saarimäki, 2017). A meta-analysis of neuroimaging studies also indicates that different basic emotion categories can be predicted across studies from activity patterns that span several intrinsic functional networks (Wager et al., 2015) suggesting that emotions elicited by non-musical stimuli arise from interactions between systems that also support processes that are not exclusively affective such as memory, attention, and action. Music-induced emotions might differ from more prototypical basic emotions by relying more heavily on sensory or aesthetic processing and consequently on more spatially localized auditory cortical representations.

### Music-induced emotions and core emotion circuits

The video experiment for mapping networks governing non-musical emotions revealed extensive and strong activation in regions involved in emotional processing including the amygdala, thalamus, NAcc, ACC and insula and midline frontal regions. Although the MVPA did not reveal consistent emotion-specific patterns outside the auditory and motor cortices, the GLM for the music-induced emotions did reveal activation in many (e.g. insula, ACC, amygdala) although not all (e.g. midline frontal regions) components of these emotion circuits.

Both the music-induced emotions (except for tenderness) and those induced by the film clips activated the ACC and insula. Previous studies indicate that ACC activity is positively associated with the intensity of pleasurable music-induced chills (Blood & Zatorre, 2001) and responds to happy music (Mitterschiffthaler et al., 2007). The insula is implicated in the processing of interoceptive feedback (Craig, 2002) and is likely involved in generating the somatic component of emotional experience (Nummenmaa et al., 2018). ACC and insula activation has consistently been coupled with autonomic arousal suggesting that these regions contribute to the physiological changes (e.g. in heart rate, blood pressure, respiration and muscle tone) that typically accompany emotions (Beissner et al., 2013). Accordingly, ACC and insula activation was particularly extensive for the high arousal emotions fear and happiness. Activation of the precuneus was also observed for most of the music-induced emotions and for the video experiment in line with previous studies showing that this region is activated by different task involving introspection (Cavanna & Trimble, 2006).

The GLM also confirmed the contribution of the amygdala in music-induced fear in accordance with human lesion studies (Gosselin et al., 2005) and some prior fMRI studies (Aubé et al., 2015; Koelsch et al., 2013, however see Bogert et al., 2016) implicating the amygdala in music-induced fear . However, liking also activated this region in line with data showing that the amygdala responds to a range of non-aversive but salient stimuli and is not exclusively involved in fear-related functions but contributes to the detection of emotional significance more generally (Pessoa, 2010; Sander et al., 2003). Accordingly, amygdala activation has previously been reported in response to both pleasant and unpleasant music compared to neutral music (Ball et al., 2007) and joyful music compared to scary music (Koelsch et al., 2006, for a meta-analysis see Koelsch, 2014).

The liking ratings also predicted activity in the hippocampus. Prior studies have associated activation in the hippocampal formation with music-induced positive emotions like tenderness (Trost et al., 2012) and joy (Mueller et al., 2011), pleasantness of musical chords (Cheung et al., 2019)), and the perception of happiness in music (Bogert et al., 2016), but also with the processing of unpleasant dissonant versions of pleasant consonant music (Koelsch et al., 2006). A cluster comprised of the hippocampus and adjacent structures was observed in a meta-analysis on the neural correlates of music-induced emotions and was proposed to mediate the stress reduction by music and music-induced positive, attachment-related emotions (Koelsch, 2014). Another study found that liked vs. disliked music activated many of the same regions whose activity correlated with the liking ratings in the current study including the caudate, thalamus, parahippocampal gyrus, anterior cingulate, superior temporal gyrus, precuneus and the cerebellum (Brattico et al., 2016). Another structure implicated in pleasure, liking and positive emotions in across studies is the NAcc (Koelsch, 2014). However, in contrast to previous studies (Blood & Zatorre, 2001; Salimpoor et al., 2013), we did not observe an association between liking and activity in the nucleus accumbens in the ventral striatum. This discrepancy might be partly due to the fact that, unlike the studies providing strongest evidence for Nacc contribution in music-induced pleasure, we did not target individuals particularly prone to experience music-induced frisson (Martínez-Molina et al., 2016; Salimpoor et al., 2013) and used unfamiliar music as stimuli (cf. Pereira et al., 2011). Furthermore, some studies reporting NAcc activity have contrasted liked music with highly unpleasant, dissonant music (Menon & Levitin, 2005; Mueller et al., 2015). In the current study, all musical stimuli received relatively high liking ratings which may have precluded the detection NAcc activity.

It is possible that the weaker limbic and paralimbic responses to music versus emotional videos simply reflect the weaker emotional potency of the musical excerpts. Alternatively, it is possible that videos contain more overlapping features driving the limbic circuits.

In sum, the GLM results indicate that music-induced emotions may engage central nodes of the core emotion circuits (cf. Koelsch, 2004) although the activation of these circuits was substantially less extensive for the music stimuli than for the videos. It is possible that the weaker limbic and paralimbic responses to music versus emotional videos simply reflect the weaker emotional potency of the musical excerpts. Alternatively, it is possible that videos contain more temporally overlapping dynamic features (e.g. facial expressions, social interaction) driving the limbic circuits (Lahnakoski et al., 2012). Irrespective of the ultimate answer to this question, activity patterns in these regions did not consistently differentiate specific music-induced emotions in the MVPA suggesting that activity in these circuits does not contain neural signatures of distinct music-induced emotions. Prior studies indicate that limbic and paralimbic regions are activated by a range of positive and negative emotions (Wager et al., 2015) perhaps because these regions govern some elementary ingredients of emotion such as encoding of valence, arousal or motivational relevance shared by all emotions. The current study suggests that, together with the emotion-specific representations in the auditory and motor cortices, these circuits give rise to subjective music-induced affective states that listeners interpret as categorically distinct basic emotions such as happiness and sadness.

### Limitations

We used unfamiliar instrumental music as stimuli to minimize the contribution of autobiographical memories and lyrics to the emotional responses. This attempt to maximize experimental control may have resulted in weaker emotional reactions than, for example, with familiar music with lyrics. Indeed, previous studies indicate that familiar music elicits stronger emotional reactions (Peretz et al., 1998) and engages the brain’s reward circuits more strongly than unfamiliar music (Pereira et al., 2011). Our self-reports are consistent with strong affective engagement (**Figure 1**), yet it is also possible that these results may be inflated due to demand characteristics. Finally, it is possible that the dimensional valence-arousal or music-specific categorical models would better describe how music-induced emotions are organized which would arguably make our analysis approach suboptimal for detecting affect-related brain responses.

## Conclusions

We conclude that music-induced emotions are represented as discrete patterns of activity in the auditory and motor cortices. These emotions consistently engage a network of auditory cortical areas (Heschl’s gyrus, planum polare, planum termporale) and regions supporting motor control, somatosensation and interoceptive processing (pre- and post-central gyri, SMA, cerebellum, ACC, insula, precuneus). Despite high statistical power, we did not find evidence for music-induced basic emotions rely strongly on the subcortical limbic system or medial frontal regions that govern basic emotions induced by biologically salient stimuli. We propose that even though listeners often interpret their subjective responses to music in terms of basic emotion categories, these feelings may only partially rely on the same neural machinery as prototypical basic emotion with adaptive significance.

