## Supplementary Material for "Decoding music-evoked emotions in the auditory and motor cortex"

#### Emotion ratings

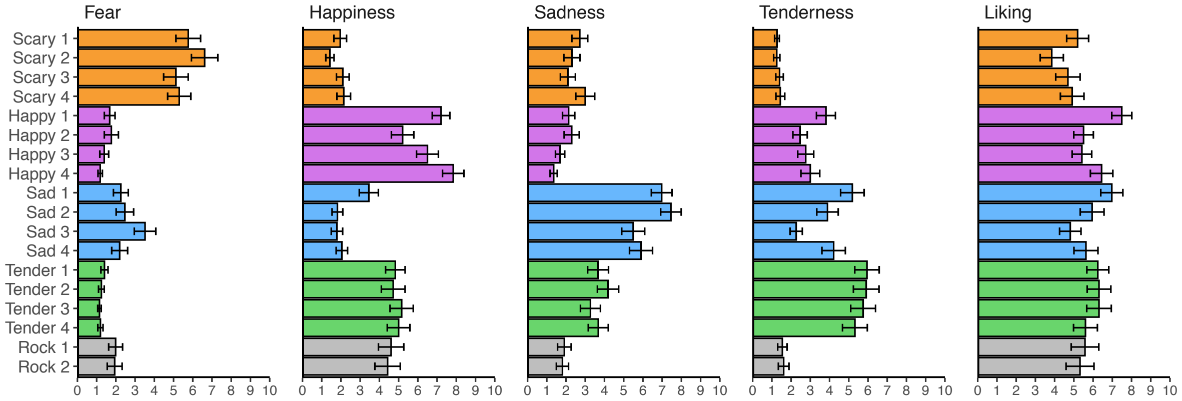


**Supplementray Figure 1.** Mean ratings for the intensity of each emotion for each musical excerpt. Error bars show standard error of mean

**Fear.** The was significant main effect of song category (scary vs. happy vs. sad vs. tender vs. rock) on the fear ratings ($F(4,360) = 152.297$, $p < .001$). Pair-wise t-test showed that the scary song were rated higher for fear than the other song: scary > happy: $t(90) = 15.386$, $p < .001$; scary > sad: $t(90) = 14.009$, $p < .001$; scary > tender: $t(90) = 16.086$, $p < .001$; scary > rock: $t(90) = 13.226$, $p < .001$. **Happiness.** Main effect of Category on happiness ratings: $F(4,360) = 123.495$, $p < .001$. Pair-wise t-test : happy > scary: $t(90) = 22.067$, $p < .001$; happy > sad: $t(90) = 20.192$, $p < .001$; happy > tender: $t(90) = 10.55$, $p < .001$; happy > rock: $t(90) = 7.168$, $p < .001$). **Sadness.** Main effect of Category on sadness ratings: $F(4,360) = 200.556$, $p < .001$. Pair-wise t-test : sad > scary: $t(90) = 21.442$, $p < .001$; sad > happy: $t(90) = 21.175$, $p < .001$; sad > tender: $t(90) = 12.437$, $p < .001$; sad > rock: $t(90) = 19.85$, $p < .001$). **Tenderness.** Main effect of Category on tenderness ratings: $F(4,360) = 165.797$, $p < .001$. Pair-wise t-test : tender > scary: $t(90) = 17.9$, $p < .001$; tender > happy: $t(90) = 11.046$, $p < .001$; tender > sad: $t(90) = 9.002$, $p < .001$; tender > rock: $t(90) = 15.517$, $p < .001$). **Liking.** Main effect of Category on liking ratings: $F(4,360) = 10.999$, $p < .001$. Pair-wise t-test : scary > happy: $t(90) = 7.929$, $p < .001$; scary > sad: $t(90) = 5.871$, $p < .001$; scary > tender: $t(90) = 5.376$, $p < .001$; scary > rock: $t(90) = 2.202$, $p < .05$); happy > rock: $t(90) = 2.707$, $p < .01$).

### Supplementary material

#### Main effect of Music vs Silence

The contrast between music vs. silence (**Supplementary Figure 1**) revealed subcortical activity bilaterally in the brainstem, thalamus, putamen, caudate, and the left pallidum. Activity was also observed in the hippocampus, parahippocampal gurys, amygdala and the insula. There was bilateral activity across the somato-motor regions in the precentral and postcentral gyri, the supplementary motor area (SMA) and the cerebellum. Occipital activity was also observed in the precuneus. Frontal activity was seen in the Inferior, middle and superior frontal gyri and the frontal pole. The cingulate gyrus showed activity in the anterior and posterior portions. Activity was also seen across the occipital cortex, fusiform gyrus, and lingual gyrus extending into the inferior temporal cortex. Finally, there was extensive activation of auditory cortical regions encompassing Heschl's gyrus, superior temporal gyrus, planum temporale and planum polare.

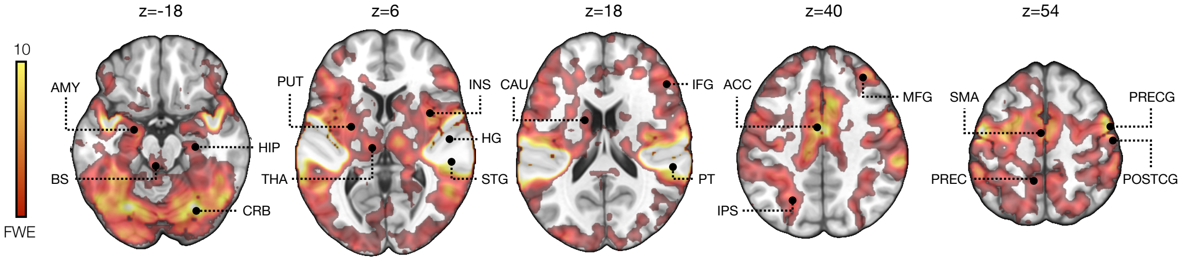


**Supplementary Figure 2.** Brain regions showing greater responses to music versus silence thresholded at $p < .05$ and FWE corrected at cluster level. ACC = Anterior Cingulate, AMY = Amygdala, CAU = Caudate, CRB = Cerebellum, FP = Frontal Pole, HC = Hippocampus, HG = Heschl's Gyrus, IFG = Inferior Frontal Gyrus, INS = Insula, PAL = Pallidum, PRECG = Precentral gyrus, PRECUN = Precuneus, SMA = Supplementary Motor Area, STG = Superior Temporal Gyrus, THA = Thalamus. The color bar indicates T-value.

#### Main effect of Music vs Control stimuli

The contrast between music vs. control stimuli (**Supplementary Figure 2**) revealed subcortical activity in the brainstem, left thalamus, right caudate, and the left pallidum. Activity was also observed in the hippocampus, parahippocampal gurys, insula, precentral and postcentral gyri, supplementary motor area, and the cerebellum. The cingulate gyrus showed activity in the anterior and posterior portions. Occipital activity was observed in the precuneus, frontal activity was observed in the middle frontal gyrus and the frontal pole, occipital activity was observed in the fusiform gyrus, and lingual gyrus, and temporal activity was observed in the auditory cortical regions.

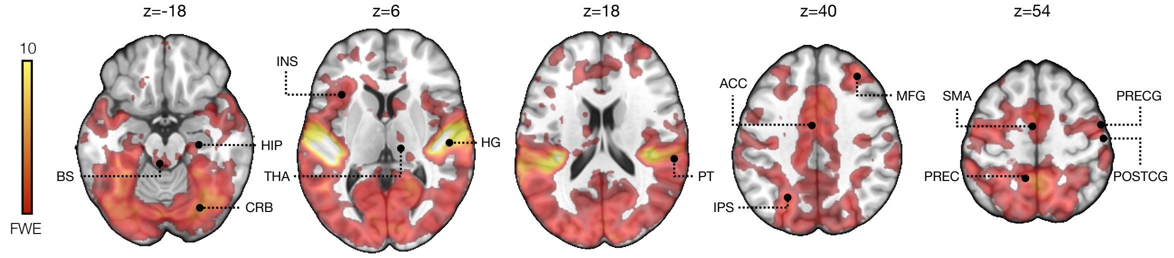

### Supplementary material

**Supplementary Figure 3.** Brain regions showing greater responses to music versus control stimuli thresholded at $p < .05$ and FWE corrected at cluster level. ACC = Anterior Cingulate, CRB = Cerebellum, HC = Hippocampus, HG = Heschl's Gyrus, INS = Insula, PRECG = Precentral gyrus, PRECUN = Precuneus, SMA = Supplementary Motor Area. The color bar indicates T-value.

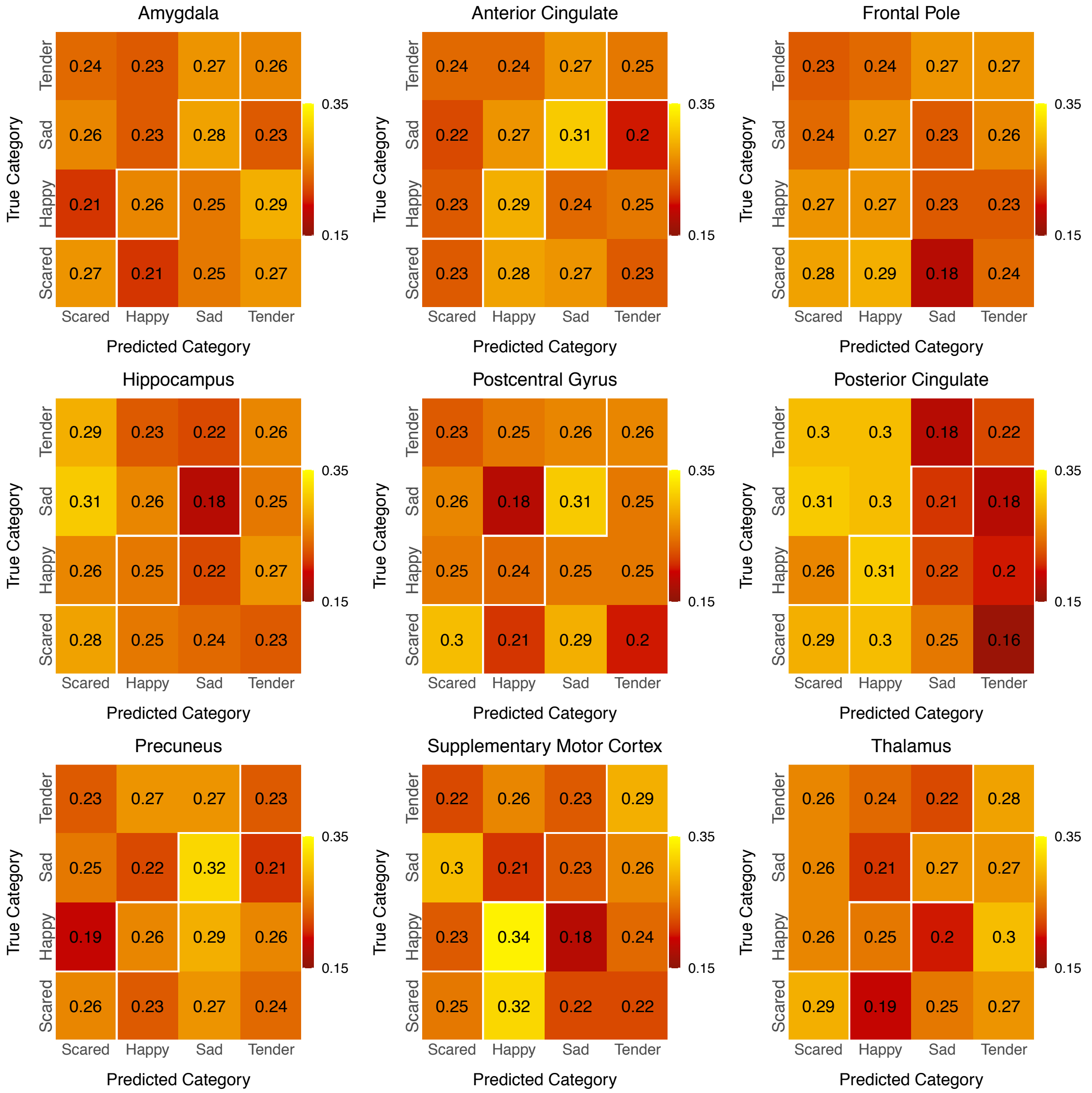


**Supplementary Figure 4.** Confusion matrices for ROIs where the classification accuracy did not reach significance for all emotions. The numbers indicate the proportion of true positives (diagonal) and false negatives within each emotion category.

### Supplementary material

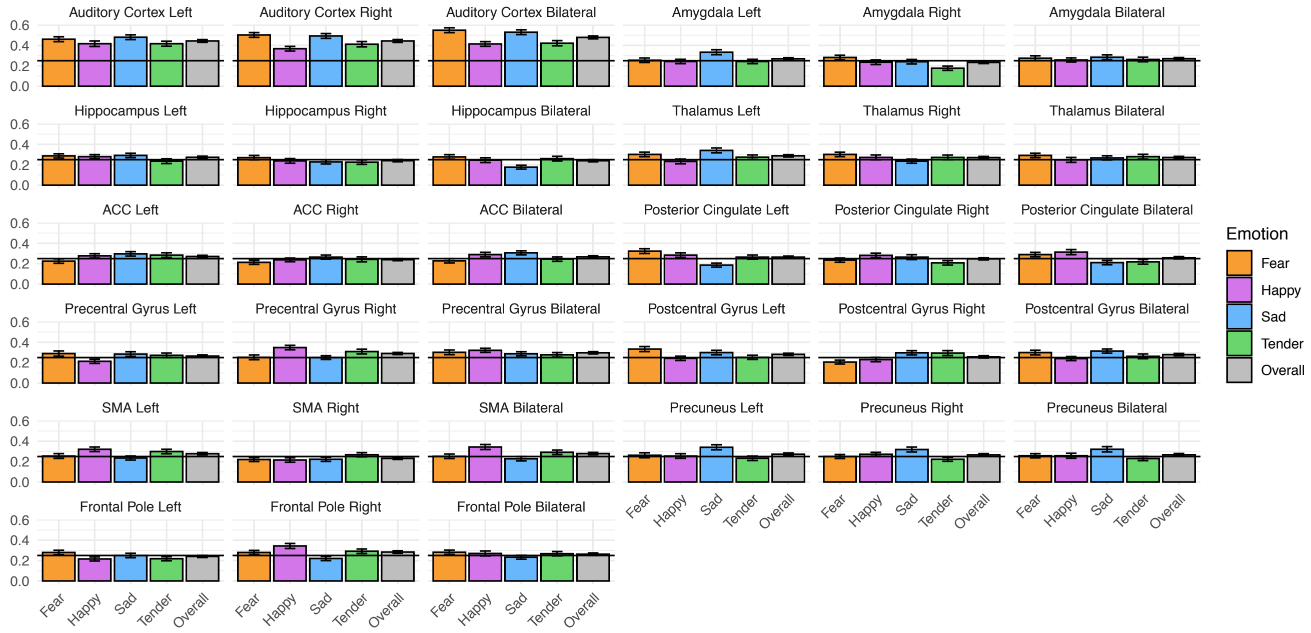


**Supplementary Figure 5.** Emotion-wise classification accuracies for the left and hemisphere and bilateral regional MVPA.
